# Depth-targeted intracortical microstroke by two-photon photothrombosis in rodent brain

**DOI:** 10.1101/2021.11.10.466928

**Authors:** Masahiro Fukuda, Takayoshi Matsumura, Toshio Suda, Hajime Hirase

## Abstract

**Significance:** Photothrombosis is a widely used model of ischemic stroke in rodent experiments. In the photothromboris model, the photosensitizer Rose Bengal is systemically introduced to the blood stream and activated by green light to induce aggregation of platelets that eventually cause vessel occlusion. Since the activation of Rose Bengal is a one-photon phenomenon and the molecules in the illuminated area (light path) are subject to excitation, targeting of thrombosis is unspecific especially in the depth dimension. We have developed a photothrombosis protocol that can target a single vessel in the cortical parenchyma by two-photon excitation.

**Aim:** We aim to induce a thrombotic stroke in the cortical parenchyma by two-photon activation of Rose Bengal so that we confine photothrombosis within a vessel of a target depth.

**Approach:** FITC-dextran is injected into the blood stream to visualize the cerebral blood flow in anesthetized adult mice with a cranial window. After a target vessel is chosen by two-photon imaging (950 nm), Rose Bengal is injected into the blood stream. The scanning wavelength is changed to 720 nm and photothrombosis was induced by scanning the target vessel.

**Results:** Two-photon depth-targeted single vessel photothrombosis was achieved with a success rate of 84.9±1.7% within 80 s. Attempts without Rose Bengal (i.e., only with FITC) did not result in photothrombosis at the excitation wavelength of 720 nm.

**Conclusions:** We described a protocol that achieves depth-targeted single vessel photothrombosis by two-photon excitation. Simultaneous imaging of blood flow in the targeted vessel using FITC dextran enabled the confirmation of vessel occlusion and prevention of excess irradiation that possibly induces unintended photodamage.

## 1 Introduction

Ischemic strokes occur at various levels of cerebral circulation ranging from massive middle cerebral artery occlusion to transient ministrokes that occur at single arterioles. Ischemia at the stroked sites causes neuronal death or irreversible damage forming an infarct due to the lack of energy substrates. Silent strokes typically occur in small arterioles and cause small-sized infarcts resulting in little expressive symptoms. These so-called microstrokes occur in the cerebral parenchyma and remain undetected until multiple occurrences of microstroke over a period integrate to develop to motor and/or cognitive deficits ^1^.

Several models have been proposed to generate local embolic or thrombotic stroke in experimental animals ^2–4^. For instance, hundreds of microspheres of diameter 20-50 µm have been perfused into the upstream of cerebral circulation ^5^, e.g., the internal carotid artery, to induce embolic ischemic stroke in the downstream arterioles. Microsphere infusion imposes a fair amount of surgical burdens and inherently forms infarcts in multiple places. Since the locations of infarcts are somewhat arbitrary and sometimes ectopic, this method is not optimal for observing the impact of single-vessel occlusion by microscopic *in vivo* imaging.

Induction of thrombosis by optic means, photothrombosis ^6^, has been widely utilized by the stroke research community owing to the technique’s reproducibility and flexible target region selection. Photothrombosis is performed by introducing the photo-sensitizer Rose Bengal to the blood and exciting it by green light in the targeted area of the brain surface. Rose Bengal excitation produces reactive oxygen species (ROS), which damages epithelial cells and promotes aggregation of platelet on the vessel wall that eventually occludes the vessel. Photothrombosis has the advantage of target area selection over other methods as the target region can be specified by excitation light illumination. However, due to the nature of one-photon reaction, photothrombosis is not suited for targeting specific vessels inside the brain tissue because the superficial vessels including capillaries are also subjected to photothrombosis with even higher intensity light. Furthermore, excess exposure of high-intensity excitation light on the mouse cerebral cortex can penetrate into deeper structures like the hippocampus and striatum and occlude off-target areas, making the interpretation of subsequent outcome unclear. Moreover, spot illumination of thrombotic target area does not distinguish between arteries and veins.

Two-photon microscopy has been utilized widely in bio-imaging due to its superior depth-penetration in the tissue and focus-specific fluorescence excitation ^7^. Taking advantage of this feature, multiphoton photothrombosis has been reported very recently by a group that employed fix-point irradiation ^8^. Here we introduce an alternative technique for depth-targeted photothrombosis (DTPT) that can readily be used on a standard two-photon microscope equipped with a resonant scanner. Using this method, we demonstrate photothrombosis in a single vessel or a group of vessels at a targeted depth of up to 300 µm. Remarkably, we show that photothrombosis is induced within 40 sec of laser illumination at the cortical depth of 300 µm without unwanted occlusions above and below the target. These features are favorable for prospective microstroke research since inducing occlusion in individual targeted vessels in the brain parenchyma is considered to mimic silent stroke.

## 2 Materials and Methods

### Subjects and surgery

Adult C57BL/6 mice (2-6 months old, 20–30 g) of either sex were used. Mice were deeply anesthetized with isoflurane (5% for induction, 1.5% for maintenance). A craniotomy of ∼3 mm diameter was performed above the somatosensory cortex while the dura was left intact. Thereafter, a glass coverslip was placed and fixed with dental cement. A metallic head frame that fits to MAG-3 head-stager holder (Narishige, Japan) was attached to the skull, and the remaining exposed skull was covered with dental cement.

The procedures involving animal care, surgery and sample preparation were approved by the Danish Animal Experiments Inspectorate and the Animal Care and Use committee of Kumamoto University, Kumamoto, Japan. The animal study was overseen by the University of Copenhagen Institutional Animal Care and Use Committee.

### In vivo two-photon depth-targeted photothrombosis (DTPT)

Two-photon imaging and depth-targeted photothrombosis (DTPT) was performed in anesthetized mice, using a Bergamo scope (Thorlabs) equipped with a resonant scanner (8 kHz), a piezo Z drive, and a 25× 1.1 NA water immersion lens (Nikon, CFI75 Apochromat 25XC W 1300), and a Mai Tai eHP DeepSee laser (Spectra-Physics). Emission light was separated by a dichroic mirror (562 nm, Chroma) with band-pass filters 525/50 nm and 607/70 nm (both from Chroma) for the green and red channels, respectively. The microscope was operated using the ThorImage LS software version 4.0.

DTPT was carried out as follows. First, to maximize the excitation efficiency, we adjusted the head-stage mount angle so that the craniotomy plane becomes parallel to the objective lens plane using the goniometer incorporated to the MAG-3 head-stage holder ^9^. Second, 300 µL of 20 mg/mL FITC-150k dextran (Sigma-Aldrich, MO, USA) was injected intravenously (i.v.) via the retro-orbital route to label blood vessels. After identifying the target blood vessel using 950 nm excitation light, 100 µL of the photo-sensitizer Rose Bengal (Sigma-Aldrich, MO, USA) was injected (i.v., Rose Bengal concentration: 20 mg/mL in phosphate buffered saline) ^10^, and injected every 20 min to maintain the RB concentration in the blood (up to three times). The excitation light wavelength was then changed to 720 nm, and the focal depth was readjusted to the target vessel’s lumen using the least laser power possible (2.9 mW). Next, the scanned area was restricted by increasing the digital magnification so to include only the targeted vessel. To start photothrombosis, the laser intensity was set to 31 mW at the exit of the objective lens, and the selected area was scanned at a high frame rate (30–60 Hz, depending on the scanned area) while monitoring acquired images in real time. If equipped, the far-red detection channel should be switched off to protect PMT. Scanning was terminated when blood clots were formed.

### Statistics

Comparisons of two sample means were assessed by t-test. Error bars of bar graphs represent the standard error of the mean. Box plots indicate the medians and 25th & 75th percentiles of the sample data. The whiskers of a box plot represent the shorter of the data range or the outlier limit. The outlier limits are defined as *q3 + 1*.*5 × (q3 – q1)* and *q1 – 1*.*5 × (q3 – q1)*, where *q1* and *q3* are the 25th and 75th percentiles of the sample data.

## 3 Results

### Repetitive fast scans in the lumen induces 2-photon photothrombosis

To undertake the induction of photothrombosis by an ultra-short pulse infrared laser, we followed a recent paper that reported an enhanced multiphoton excitation of Rose Bengal for wavelengths < 750 nm ^11^. First, we visualized the cerebral vasculature by labeling the serum with FITC-dextran via i.v. injection and imaging through the cranial widow by a two-photon microscope in an anesthetized mouse (Fig. 1(a)). A wavelength of 950 nm was used to excite FITC. Cerebral arterioles were identified by tracking the downstream of a descending blood flow in the penetrating arteries. A vessel of diameter 4–20 µm was chosen as a target and placed at the middle of the view field by adjusting the microscope stage. The scanned area was restricted to the target vessel by zooming into the vessel (Fig. 1(b)). After injecting Rose Bengal i.v., the wavelength was changed to 720 nm and the scanned depth (the Z-axis focus) was readjusted to scan inside the lumen with a minimum laser intensity with a frame rate at 30 or 60 Hz. After setting the target, the laser power was set to 31 mW (at the exit of the objective lens). The PMT voltage and amplifier gain was set to low values to allow for simultaneous photothrombosis and vessel blood flow monitoring.

**Fig. 1.**
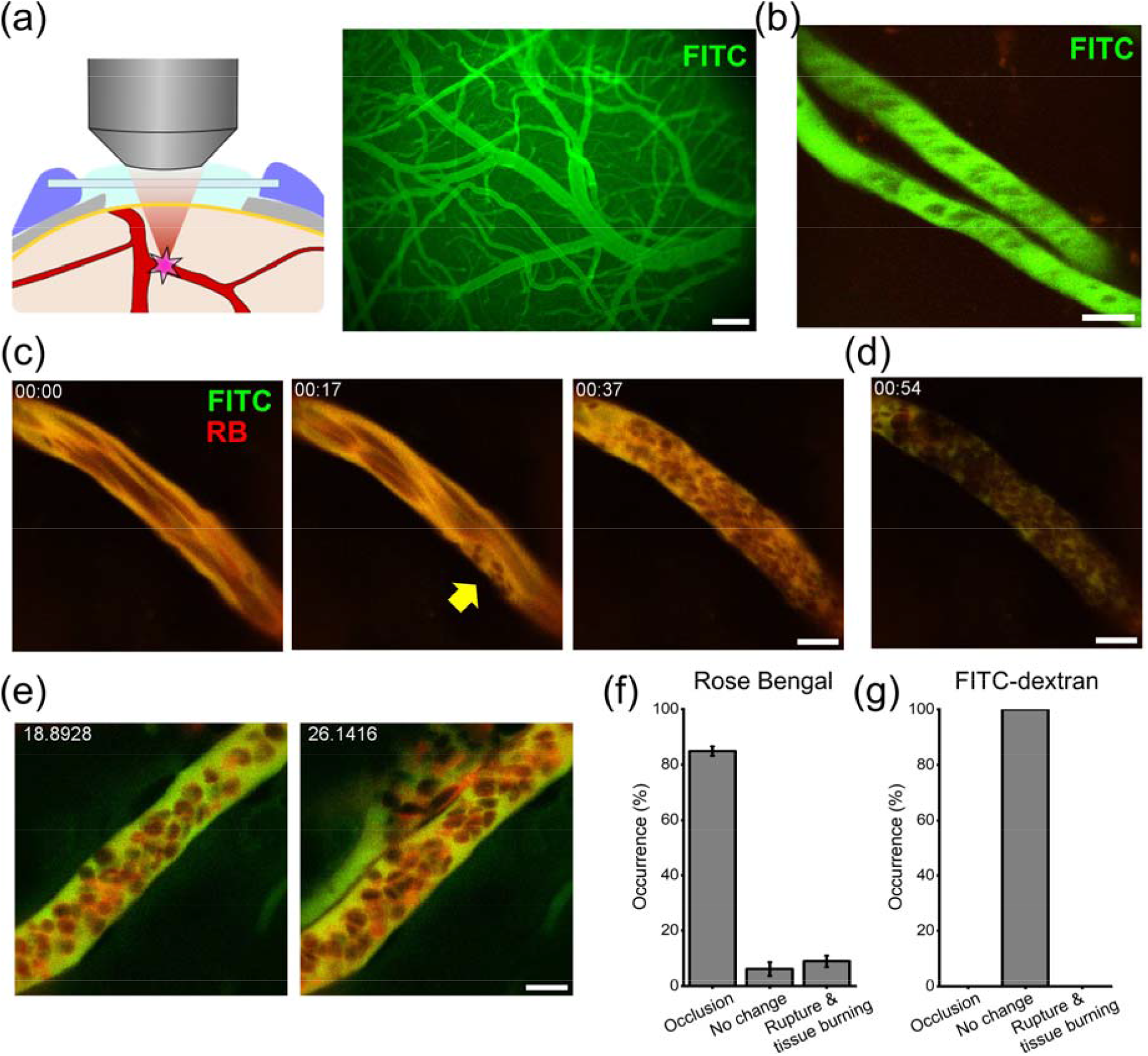
*In vivo* two-photon depth-targeted photothrombosis (DTPT): (a) Schematic illustration of DTPT. After visualization of blood vessels with FITC-dextran, a blood vessel containing Rose Bengal is targeted by 2-photon excitation. (b) Blood vessels to be targeted were chosen using FITC-dextran fluorescence. (c) Following Rose Bengal injection, the excitation wavelength was switched to 720 nm. Imaging of the targeted blood vessel with 720 nm wavelength was performed until coagulation was formed. (d) Completion of occlusion could be distinguished by photobleaching of Rose Bengal and FITC. (e) In a small number of cases, blood vessel rupture or tissue burning was observed. (f) The success rate of DTPT was 84.9±1.7%, while 6.1±2.5% of the cases were not occluded (No change) and 9.0±2.1% of the cases had tissue burning/rupture (occlusion: 84/99, no change: 6/99, tissue burning/rupture 9/99 vessels, 4 mice). (g) Controls with FITC-150k dextran only did not show occlusion or tissue burning/rupture in 150 sec (103 blood vessels, 3 mice). Scale bars: (a) 200 µm; (b),(c),(d),(e): 10 µm.

**Video 1.**
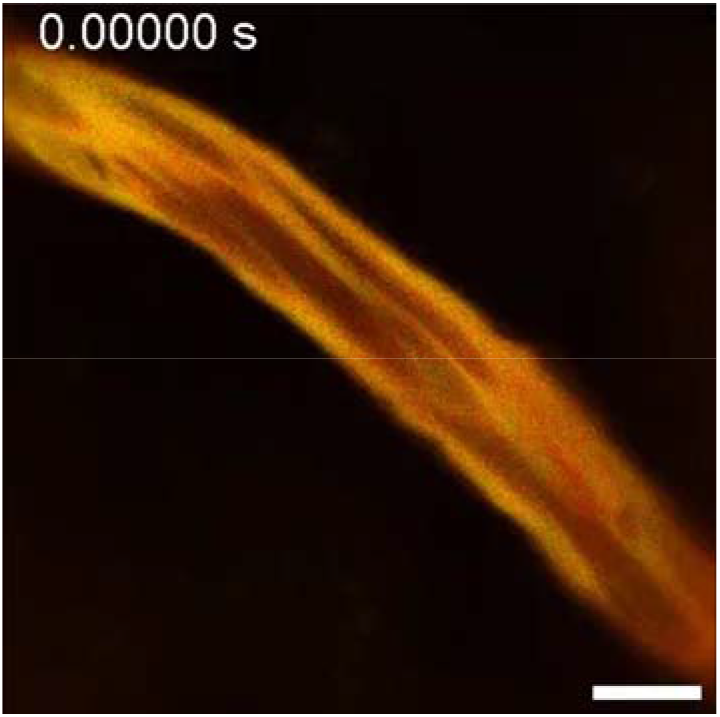
Occlusion of a single cortical vessel by DTPT. Scale bar 10 µm. (MP4, 7MB)

The blood flow of the subjected vessel was recognized from the suboptimal fluorescence of Rose Bengal and FITC as acquired images are updated on the screen (Fig. 1(c), left). Initially, platelets and red blood cells started to attach to the blood vessel wall (Fig. 1(c), middle). Thereafter, red blood cells (RBCs) made a clot, and the blood flow was gradually compromised (Fig. 1(c), right; Video 1). Occlusion of the vessel was observed by the appearance of stalled RBCs in the image. In addition, the blood-contained dyes quicky became bleached and after complete occlusion, which can be used as the sign of occlusion (Fig 1(d)). Occlusion was induced within 80 s of the 720 nm irradiation in the majority of cases (Fig. 1(f), 84/99 vessels, 4 mice), though we have observed failure of occlusion or rupture/tissue burning in a small number of attempts (Fig. 1(f)). In contrast, FITC-only control displayed neither occlusion nor attachment of platelets to the vessel walls even when they were irradiated for up to 150 sec (Fig. 1(g)).

We did not find that the diameter of blood vessels affected the success of occlusion formation (Occlusion: 11.58 ± 0.43 µm (n=84), No change: 8.27 ± 1.57 µm (n=6), P=0.069) (Fig. 2(a)). We find that the success rate of occlusion formation by DTPT is affected by the time since Rose Bengal injection. The unsuccessful attempts occurred when significantly longer time have passed after Rose Bengal injection than typical successful attempts (successful occlusion: 9.0±0.6 minutes (n=84), No occlusion: 13.8±1.6 minutes (n=6) since Rose Bengal injection. P=0.019) (Fig. 2(b)). These results together with FITC-only control (Fig. 1(g)) indicated that the concentration of Rose Bengal inside the blood vessel is crucial for DTPT and 720 nm excitation laser reliably excites Rose Bengal.

**Fig. 2.**
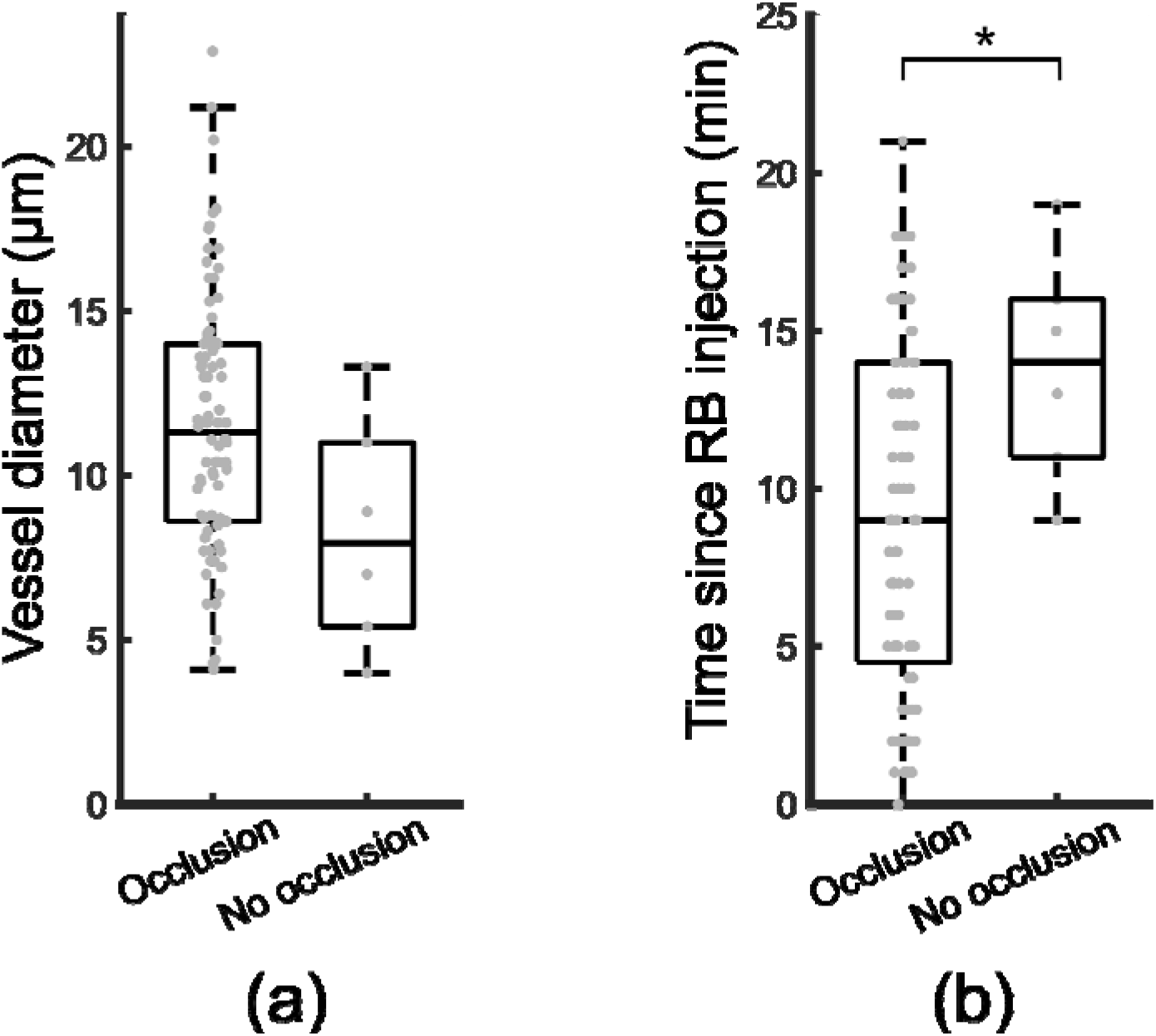
Parameters for successful DTPT: (a) Distribution of vessel diameter for successfully occluded vessels vs. not occluded vessels. (b) Distribution of time since Rose Bengal (RB) injection for successfully occluded vessels vs. not occluded vessels. * P < 0.05.

### DTPT occlusion is confined to the targeted vessels

While the DTPT attempts we described thus far have been done in target planes 30–50 µm from the pial surface, we next performed DTPT targeting the blood vessel (φ=6.5 µm) located at the depth of 300 µm. In this case, we used the laser power ∼300 mW for the 720 nm irradiation.

Accordingly, and the blood clot was formed within 40 sec (Fig. 3(a) and (b), Video 2). The clot formed was clearly visible with 950 nm excitation (Figure 3(c), arrow). The utility of DTPT becomes powerful when off-focus vessels are not occluded. To ensure that photothrombosis was confined to the targeted vessel, we took volumetric images and examined whether there are any blood clots in the capillaries 100, 200 and 300 µm above and 100 µm below the target plane shortly after DTPT (within 5 min) at a wavelength of 950 nm. As shown in a representative image in the result, we saw no obvious clots in the off-target planes (Fig. 3(d) and Video 3). To examine the effect of selective occlusion, we observed the area 3 days later. We found that the peripheral blood vessels of the targeted plane were also occluded, and extravasation of FITC around the occuluded blood vessels were discernible (Fig. 3(e)).

**Fig. 3.**
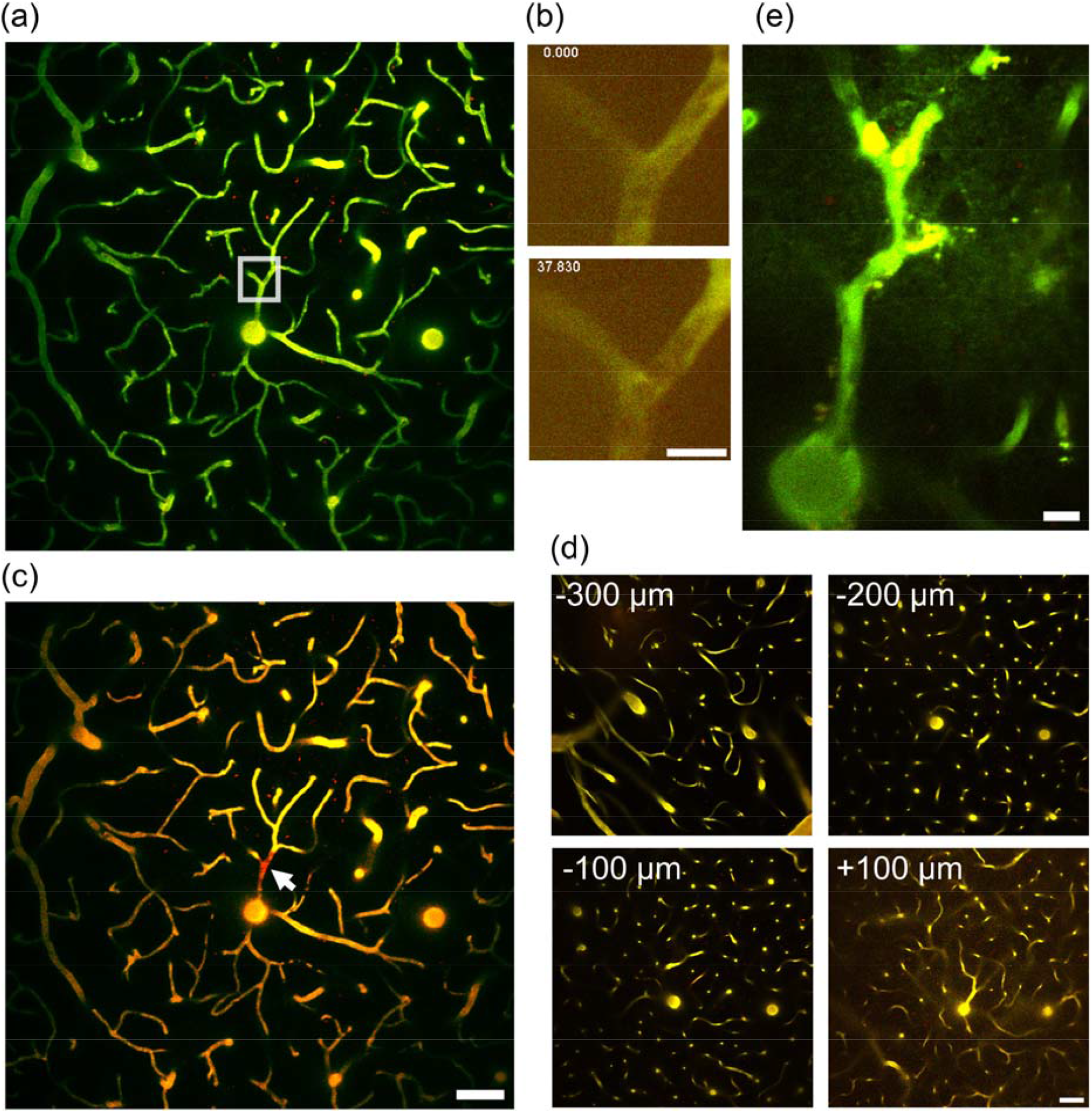
Off-target effect of DTPT is negligible: (a) The blood vessel located at 300 µm from the pia was targeted. (b) Occlusion was formed in 40 sec using 300 mW 720 nm excitation laser. (c) After DTPT, the excitation was set to 950 nm, and coagulation was visualized. (d) Even using 300 mW excitation for DTPT, there was no occlusion/coagulation in areas 300, 200, 100 µm above and 100 µm below the target. (e) Extravasation of FITC-150k dextran and occlusion of peripheral blood vessels were observed three days after DTPT. Scale bars: (a),(c),(d) 50 µm; (b),(e) 10 µm.

**Video 2.**
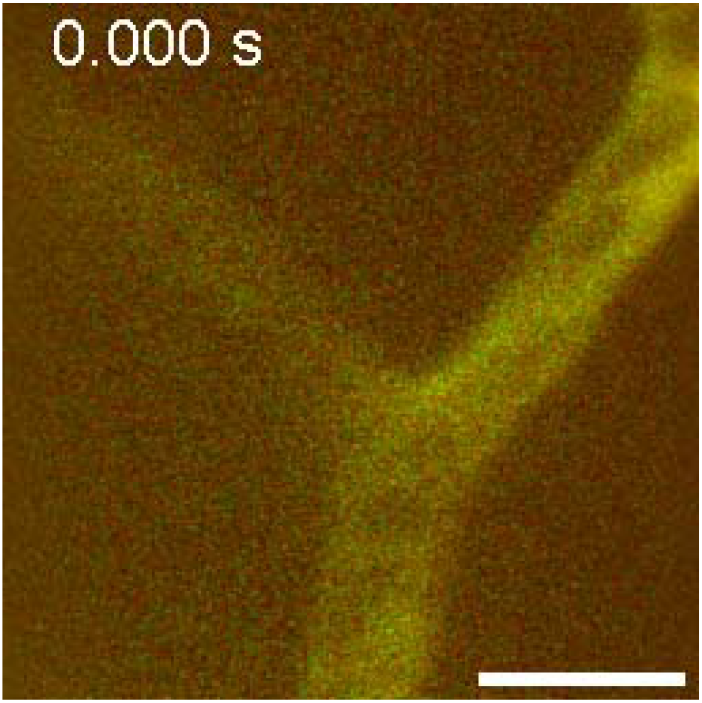
Occlusion of the vessel corresponding to Fig. 3(b). Scale bar 10 µm. (MP4, 5 MB)

**Video 3.**
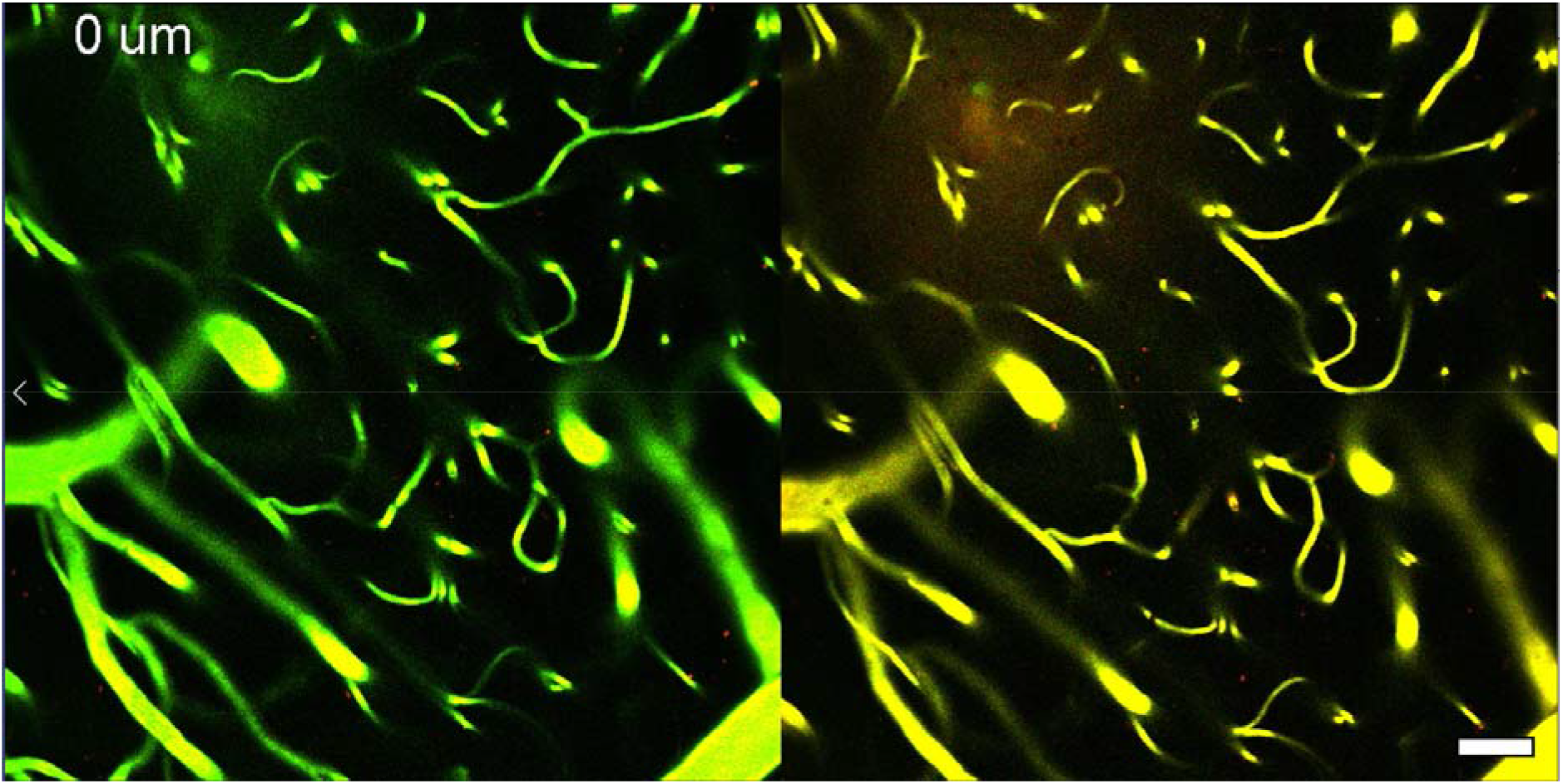
Depth stack of the cortical vasculature around the DTPT target site. Scale bar 10 µm. (MP4, 12 MB)

## 4 Discussion

Photothrombosis has been introduced for over three decades ^6^ and remains widely used for targeted induction of ischemic stroke owing to its superior reproducibility. Despite the widespread utility, photothrombosis has not been utilized for induction of microstrokes targeted specifically at a depth of the cerebral cortex until very recently. This is because conventional photothrombosis relies on one-photon excitation of Rose Bengal: ROS is produced wherever the green photon hits Rose Bengal, hence the formation of thrombosis is biased towards superficial vessels. On the other hand, multiphoton excitation occurs only at the focal spot avoiding the production of ROS production outside the focal plane and thereby enables depth-specific targeting. Indeed, Delafontaine-Martel and colleagues have demonstrated that multiphoton occlusion is feasible by two-photon or three-photon excitation. In their study, two-photon photothrombosis was achievable in depths up to 200 µm from the pial surface, and significantly longer irradiation time (up to 300 seconds) was needed with an excitation wavelength of 1000 nm ^8^.

In the current work, we further confirm that two-photon photothrombosis is feasible in the mouse brain. Notably, we find that less than 80-s irradiation is sufficient to induce microvessel occlusion at a success rate > 80%. This accelerated two-photon photothrombosis was possible with a significantly shorter wavelength of 720 nm. The one-photon absorption spectrum of Rose Bengal has a broad peak at around 560 nm, and green light of range 530–580 nm is typically used for Rose Bengal activation. Recently, Campaign and Knox have measured the two-photon excitation spectrum of Rose Bengal ^11^. According to this study, the two-photon-excited fluorescence increases by approximately three-fold as the excitation wavelength shifts from 780 nm to 720 nm. While the fluorescence emission does not necessarily correlate to ROS production, the demonstration of elevated molecule excitability at 720 nm prompts a future study to characterize ROS production around this wavelength. Another methodological point we consider important for the induction of efficient photothrombosis is that we scan the lumen of the target vessel and avoid irradiating vessel walls (endothelial cells) as much as possible. This procedure, combined with simultaneous irradiation and imaging, is designed for ensuring excitation of Rose Bengal without damaging the vessel integrity. For that matter, it is imperative to perform DTPT shortly after Rose Bengal administration so that the plasma retains sufficient Rose Bengal levels.

Although not attempted in this study, the shortened time of photothrombosis induction makes it likely that this method is applicable to awake mice, which eliminates confounding factors associated with anesthetics ^12^ and hence mimics human strokes more closely. A previous study has documented formation of single-vessel clot using a kilohertz pulsed laser of pulse energy a few orders of magnitude more intense than a typical laser used for two-photon imaging ^13^, however, this method requires intricate extra optics setup to produce the high-energy pulses. Our method works on a standard two-photon microscope without any hardware modifications hence appeals for simplicity and versatility.

Single-vessel photothrombosis has been performed on surface vessels ^14–18^, but the supposed off-target effects on deeper parenchymal vessels have not been thoroughly addressed. A clever application of one-photon photothrombosis is targeting penetrating vessels ^19^, thereby the partially circumventing the issue of depth-axis-wide activation of Rose Bengal. DTPT offers a distinct advantage over the conventional green light-induced photothrombosis by using infrared light that penetrates deeper into the parenchyma. Hence, DTPT should be an effective means to evaluate the theoretical prediction of microstroke outcomes in various microcirculation configurations ^20^. Future applications of the DTPT include occlusion of subcortical regions such as the hippocampus, where two-photon or three-photon imaging from the cortical surface is possible ^9,21–23^. Our preliminary attempts suggest that this is feasible although a thorough histological investigation of the cortical areas in the passage of the infrared beam should be performed to confirm target-specific induction of thrombotic stroke.

Another future utility of DTPT is to investigate the impact of single microvessel occlusion on the surrounding neuropil activity and inflammation. Astrocytes, a major glia subtype, have been reported to elicit aberrant activities after wide field photothrombosis ^24,25^. Chronic imaging Ca^2+^ activity or morphology will be a great resource to understand how neuropils normalizes after the mild ischemic insult or how angiogenesis possibly occurs. Of note, we have recently described that adrenergic receptor antagonism accelerates neuronal activity recovery after wide field photothrombotic stroke ^25^ or cortical spreading depression/depolarization ^26^. It would be interesting to examine how adrenergic receptor antagonism affects the homeostasis of the extracellular environment around a singly occluded vessel in light of the adrenergic receptor-dependent glymphatic system ^27,28^.

## Acknowledgments

MF is supported by JSPS Grant-in-Aid for Scientific Research (S) (JP18H05284 to T. Suda) and JSPS Grant-in-Aid for Challenging Research (Exploratory) (JP21K19514 to T. Matsumura). HH is supported by the Novo Nordisk Foundation (NNFOC0058058), Danmarks Frie Forskningsfond (0134-00107B), the Lundbeck Foundation.

## Code, Data, and Materials Availability

All materials used in the current procedure are available commercially. ThorImage LS is proprietary image acquisition software for Thorlab’s multiphoton microscopes.

